# Foxd1 dependent induction of temporal retinal character is required for visual function

**DOI:** 10.1101/2022.05.12.491645

**Authors:** María Hernández-Bejarano, Gaia Gestri, Clinton Monfries, Lisa Tucker, Elena I. Dragomir, Isaac H. Bianco, Paola Bovolenta, Stephen W. Wilson, Florencia Cavodeassi

**Author notes:** Correspondence: Florencia Cavodeassi, Gaia Gestri, Stephen W Wilson. Rights retention statement: This research was funded in part, by the Wellcome Trust [Seed Award in Sciences 213928/Z/18/Z to FC; Investigator Award 104682/Z/14/Z to SWW; Senior Research Fellowship 220273/Z/20/Z to IHB]. For the purpose of open access, the authors have applied a CC BY public copyright licence to any Author Accepted Manuscript version arising from this submission.

## Abstract

Appropriate patterning of the retina during embryonic development is assumed to underlie the establishment of spatially localised specialisations that mediate the perception of specific visual features. For instance, in zebrafish, an area involved in high acuity vision (HAA) is thought to be present in the ventro-temporal retina. Here we show that the interplay of the transcription factor Rx3 with Fibroblast Growth Factor and Hedgehog signals, initiates and restricts *foxd1* expression to the prospective temporal retina, initiating naso-temporal regionalisation of the retina. Abrogation of FoxD1 results in the loss of temporal and expansion of nasal retinal character, and consequent absence of the HAA. These structural defects correlate with severe visual defects as assessed in optokinetic and optomotor response assays. In contrast, optokinetic responses are unaffected in the opposite condition in which nasal retinal character is lost at the expense of expanded temporal character. Our study indicates that the establishment of temporal retinal character during early retinal development is required for the specification of the HAA, and suggests a prominent role of the temporal retina in controlling specific visual functions.

**Summary statement:** This study provides a mechanistic link between eye patterning and the establishment of functionally distinct retinal regions and reveals the temporal retina preferentially controls specific aspects of visual function.

## Introduction

The position at which retinal ganglion neurons differentiate along the naso-temporal and dorso-ventral axes of the forming eye determines the location at which they innervate central targets, ensuring accurate retinotopic connectivity. Consequently, the mechanisms by which retinal neurons acquire their regional identity are critical for correct visual function. Regionally localised specialisations of the retina include the high acuity areas (HAAs) described in many diurnal organisms. In many primates and birds, this structure is called the fovea and is characterized by a morphological indentation, high density of RGCs and cone photoreceptors, absence of rod photoreceptors, specialised inner retina neuronal circuitry, and absence of vasculature (Bringmann et al., 2018; Bringmann, 2019; da Silva and Cepko, 2017; Kolb et al., 2020; Kozulin et al., 2009; Kozulin et al., 2010). In teleost fish such as the zebrafish, some features associated with the fovea are evident in the ventro-temporal region of the retina, suggesting this region is comparable to the fovea in birds and primates (Mangrum et al., 2002; Pita et al., 2015; Schmitt and Dowling, 1999; Yoshimatsu et al., 2020; Zhou et al., 2020; Zimmermann et al., 2018). During development, the human fovea shows precocious expression of neuronal differentiation genes (Hoshino et al., 2017). In the chick, DaSilva and Cepko (2017) identified low levels of retinoic acid (RA) activity and high levels of the Fibroblast Growth Factor ligand FGF8 in a highly circumscribed area of the retina, prefiguring the fovea, a pattern that is conserved in the human presumptive fovea (Cornish et al., 2004; Cornish et al., 2005; da Silva and Cepko, 2017).

HAAs are generally located in the temporal retina (Bringmann, 2019; Kolb et al., 2020; Pita et al., 2015), suggesting their specification may be dependent upon acquisition of temporal character. In the zebrafish, the subdivision of the retina in domains with nasal and temporal character is evident from early stages of optic vesicle evagination by the expression of *foxg1* in the future nasal half and *foxd1* in the future temporal half of the eye (Hernández-Bejarano et al., 2015; Picker et al., 2009). Initially, the naso-temporal subdivision is aligned with the dorso-ventral axis of the developing central nervous system. However, as eye morphogenesis progresses, the eye primordium rotates such that nasal and temporal retinae relocate to their final position, aligned with the anterior-posterior axis (Kwan et al., 2012; Picker et al., 2009).

Previous studies have shown that the establishment of nasal and temporal character requires the spatially localised activity of the Sonic-Hedgehog (Shh) and Fgf signalling pathways (Hernández-Bejarano et al., 2015; Picker and Brand, 2005; Picker et al., 2009). Shh, expressed in ventral midline structures promotes *foxd1* expression in the ventral (future temporal) half of the optic primordium. Fgfs, emanating from dorsal forebrain and adjacent tissues induce *foxg1* in the dorsal (future nasal) optic primordium and repress *foxd1* expression, contributing to its confinement ventrally. Cross-repression between FoxG1 and FoxD1 at the border between the dorsal and ventral halves of the eye primordium subsequently refines the NT subdivision (Hernández-Bejarano et al., 2015; Takahashi et al., 2009; Takahashi et al., 2003). When Shh signalling is absent, *foxd1* expression can be restored by suppressing Fgf signalling indicating that an additional factor(s) can promote *foxd1* expression independently of Shh and Fgf.

Here we elucidate the molecular mechanisms involved in establishing temporal fate during retinal development and explore how alterations to naso-temporal retinal patterning impact HAA formation and visual function. We identify the transcription factor Rx3 (Loosli et al., 2003; Loosli et al., 2001) as being required for *foxd1* expression in the forming eye. *rx3* is expressed in the eye field and is required for eye formation in all studied vertebrates (Andreazzoli et al., 1999; Andreazzoli et al., 2003; Bailey et al., 2004; Fish et al., 2014; Loosli et al., 2003; Loosli et al., 2001; Mathers et al., 1997; Voronina et al., 2004). We show that the interplay of Rx3 with Shh and Fgf activities initiates and restricts *foxd1* expression to the prospective temporal half of the evaginating optic vesicles. We further show that the establishment of a *foxd1* expression domain in the eye is linked to the formation of the HAA. Larvae lacking FoxD1 function show normal retinal neuron differentiation and lamination, but temporal retinal and HAA markers are absent, and nasal markers expand throughout the retina. In contrast, fish in which nasal retinal character is absent and temporal character is expanded, show a concomitant expansion of HAA markers. In both conditions, retinotectal projections are severely perturbed. Despite this, only fish lacking FoxD1 show significant visual impairment, which was revealed by examining optokinetic and optomotor reflex behaviours and suggests impaired perception of whole-field visual motion. Our study reveals a prominent role of the temporal retina in controlling specific aspects of visual function and provides a mechanistic link between early patterning of the eye primordium and efficient visual performance.

## Results

### 1- Rx3 provides competence for Shh to promote *foxd1* expression in the ventral portion of the optic vesicle

Our previous studies (Hernández-Bejarano et al., 2015) showed that spatially localised Shh and Fgf signalling establishes the discrete domains of *foxd1* and *foxg1* expression in the presumptive retina. However, those studies also showed that blocking the activity of both Shh and Fgf in otherwise wildtype embryos resulted in expression of *foxd1* throughout the optic vesicle, suggesting that an additional factor can promote *foxd1* expression in absence of normally critical signals.

The transcription factor Rx3 is specifically expressed in the eye field and is part of a network of regulatory proteins that confers eye fate and promotes the evagination of the optic vesicles. Lack of Rx function results in eye loss (Andreazzoli et al., 1999; Andreazzoli et al., 2003; Loosli et al., 2003; Mathers et al., 1997; Voronina et al., 2004). However, spatially localised expression of *rx3* (prospective retina marker), *nkx2*.*1* (prospective hypothalamic marker) and *emx3* (prospective telencephalic marker) is initially normal in the brain and eye-forming regions of *rx3*^*−/−*^ embryos (**Figure 1A-D**; Kennedy et al., 2004) indicating that in the mutants the eye-forming territory is still present and molecularly distinct from surrounding anterior neural structures. The presence of a prospective eye-forming domain has enabled us, in this study, to assess the effect of loss of *rx3* on expression of genes that delineate NT patterning.

**Figure 1:**
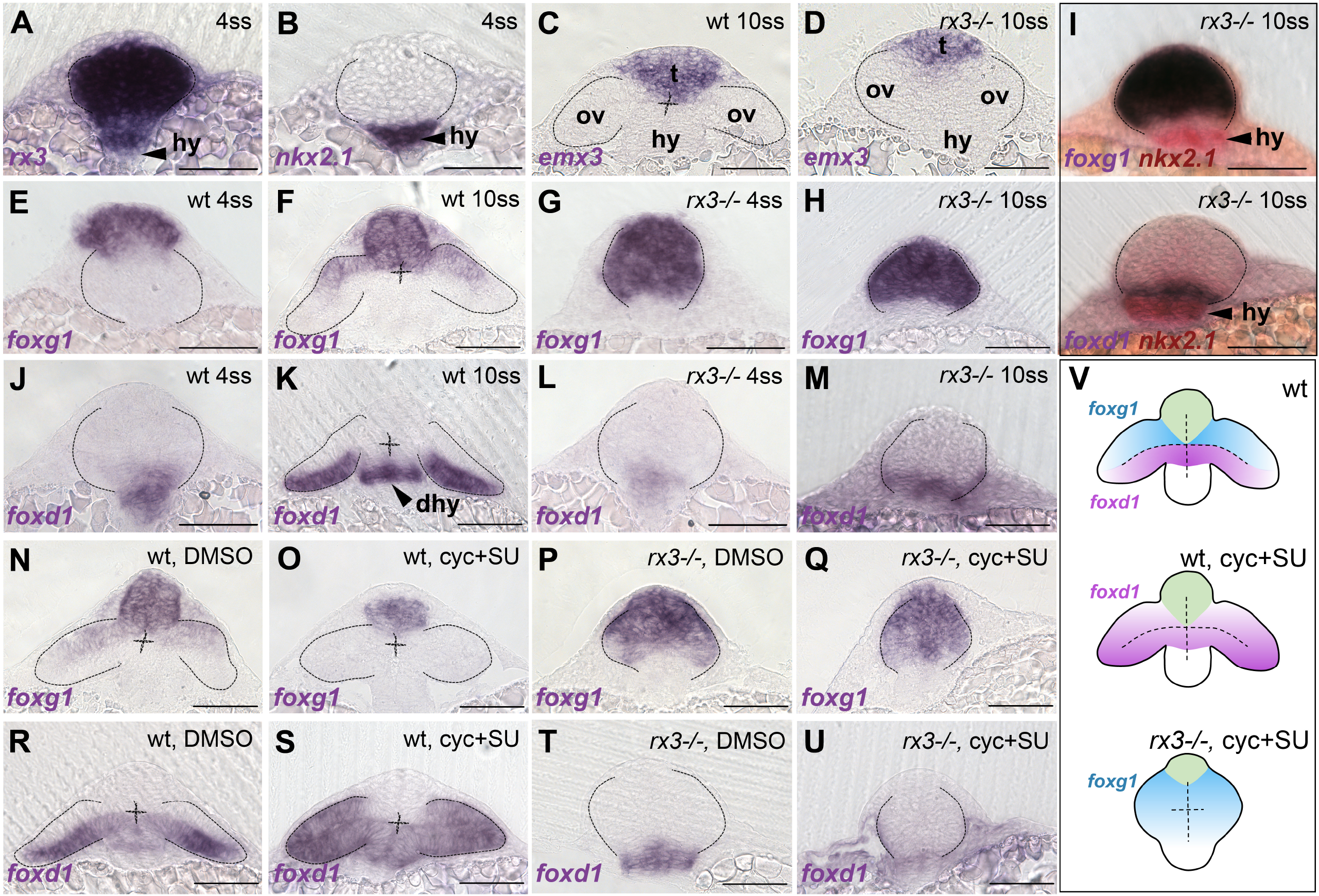
*foxd1* expression is reduced in *rx3*^*−/−*^ mutant optic vesicles and lost in *rx3*^*−/−*^ upon combined abrogation of Shh and Fgf. (A-U) Frontal sections at the level of the forming telencephalon, optic vesicles and hypothalamus, with dorsal up, genotype/treatment top right and genes analysed bottom left. (A-D) Expression of *rx3* (A) and *nkx2*.*1* (B) in 4ss wildtype/*rx3*^*−/−*^ embryos (indistinguishable at this stage with a representative embryo shown) and *emx3* in 10ss wildtype (C) and *rx3*^*−/−*^ (D) embryos highlighting the prospective telencephalic (t), hypothalamic (hy) and eye-forming (ov) domains. (E-M) Expression of *foxg1* (E-H) and *foxd1* (J-M) in 4ss (E,G,J,L) and 10ss (F,H,K,M) embryos (genotype top right). Panel (I) shows double *in situ* hybridisation of *foxd1* or f*oxg1* and *nkx2*.*1* (red) to show relationship between optic vesicles and hypothalamus. (N-U) 10ss stage embryos showing expression of *foxg1* (N-Q) or *foxd1* (R-U) in embryos treated with DMSO (N,P,R,T) or cyclopamine+SU5402 (O,Q,S,U). (V) schematic representation of the conditions in (R-U). Scale bar in this and all figures unless specified: 100μm. Dashed lines in this and all figures highlight the contour of the optic vesicles. Abbreviations: ov, optic vesicles; hy, hypothalamus; dhy, dorsal hypothalamus; t, telencephalon.

*foxg1*, expressed in the telencephalon and nasal half of the eye primordium (**Figure 1E-F**; Hernández-Bejarano et al., 2015), was expanded throughout most of the prospective eye domain in *rx3*^−/−^ mutants (**Figure 1G-H**; Stigloher et al., 2006), while *foxd1* expression was much reduced (**Figure 1J-M**). We confirmed the changes in the extent of *foxd1*/*foxg1* expression were localised to the prospective eye, by comparing their expression domains to those of *rx3* itself (which labels the eye field) and *nkx2*.*1* (**Figure 1A-B, I**). A previous transcriptomic analysis in *rx3*^−/−^ mutants suggested that *foxd1* may be a downstream target for Rx3 (Yin et al., 2014). That study, together with our observations, support a scenario in which Rx3 is required for *foxd1* expression in the forming eye and provides cells with the competence to respond to Shh in promoting the expression of *foxd1*. If this is the case, the widespread expression of *foxd1* in the optic vesicle following loss of Shh/Fgf signalling should be absent in *rx3*^*−/−*^ embryos. To address this, we first assessed whether Shh and Fgf signalling is overtly altered in *rx3*^*−/−*^ mutants.

*rx3*^*−/−*^ embryos showed normal levels of expression of genes encoding Fgf and Shh ligands and target genes in the forebrain, suggesting that levels of activity of these two pathways in *rx3*^*−/−*^ mutants are similar to wild types. The ligand encoding genes *fgf8* and *shh* showed only slight changes at the most anterior portion of the forebrain [compare the extent of *fgf8* (brackets, **Figure S1A-B**) and *shh* (arrows, **Figure S1G-H**) expression between wildtype and *rx3*^*−/−*^ embryos]. In addition, expression of *pea3* and *erm* (direct targets of the Fgf pathway) and *gli1* and *patched 2* (direct targets of the Shh pathway), was largely normal (**Figure S1C-F**,**I-L**). To abrogate Shh and Fgf activity in *rx3*^*−/−*^ embryos, we treated clutches of embryos obtained from *rx3*^*+/−*^ in-crosses simultaneously with two drugs: cyclopamine, which inhibits the Hh transducer Smoothened; and SU5042, an inhibitor of Fgf receptors (as previously described in Hernández-Bejarano et al., 2015).

In contrast to cyclopamine+SU5042-treated wildtype embryos in which *foxd1* expression is expanded (**Figure 1R-S**; Hernández-Bejarano et al., 2015), the residual expression of *foxd1* in *rx3*^*−/−*^ embryos (**Figure 1L-M,T**) was completely lost when embryos were treated with cyclopamine+SU5042 (**Figure 1T-V**; 30-50 embryos were treated and processed together per experiment; *rx3*^*−/−*^ embryos were recovered at the expected mendelian proportions and the phenotypes observed were fully penetrant). Remarkably, *foxg1*, which was lost in the optic vesicles of wildtype embryos treated with cyclopamine+SU5402 (**Figure 1N-O**), was expressed throughout the eye field of *rx3*^*−/−*^ embryos regardless of treatment (**Figure 1P-Q,V**). This may be due to the lack of the *foxg1* repressor FoxD1 in these conditions. Alternatively, it may be a consequence of the progressive adoption of telencephalic character by the eye-forming region of *rx3*^*−/−*^ mutants (Stigloher et al., 2006). Consistent with the first scenario, *foxg1* was not expressed in the prospective eye region of *rx3*^*−/−*^ embryos treated only with the Fgf inhibitor SU5402 (**Figure S2G-H**), in which there was expanded *foxd1* expression (**Figure S2E-F**).

### 2- In absence of *foxd1*, retinae fail to develop temporal character

The results above indicate a critical role for Rx3 and Shh in the induction of *foxd1* expression and the establishment of temporal character in the evaginating optic vesicle. Studies in other model organisms have shown that FoxD1 has a prominent role as determinant of retinal temporal character (Carreres et al., 2011; Hatini et al., 1994; Herrera et al., 2004; Takahashi et al., 2009; Takahashi et al., 2003). To further explore the role of *foxd1* in promoting temporal character, we generated a zebrafish loss of function mutant using CRISPR-Cas9 technology (Prykhozhij et al., 2017; see Materials and Methods). A 10 base-pair deletion just prior to the sequence encoding the FoxD1 DNA binding domain led to a frame-shift and a stop codon in position 70 of the protein product (*foxd1*^*cbm16*^; **Figure 2A-B**). This mutation likely produces a non-functional form of the protein, truncated just prior to the DNA binding site.

**Figure 2:**
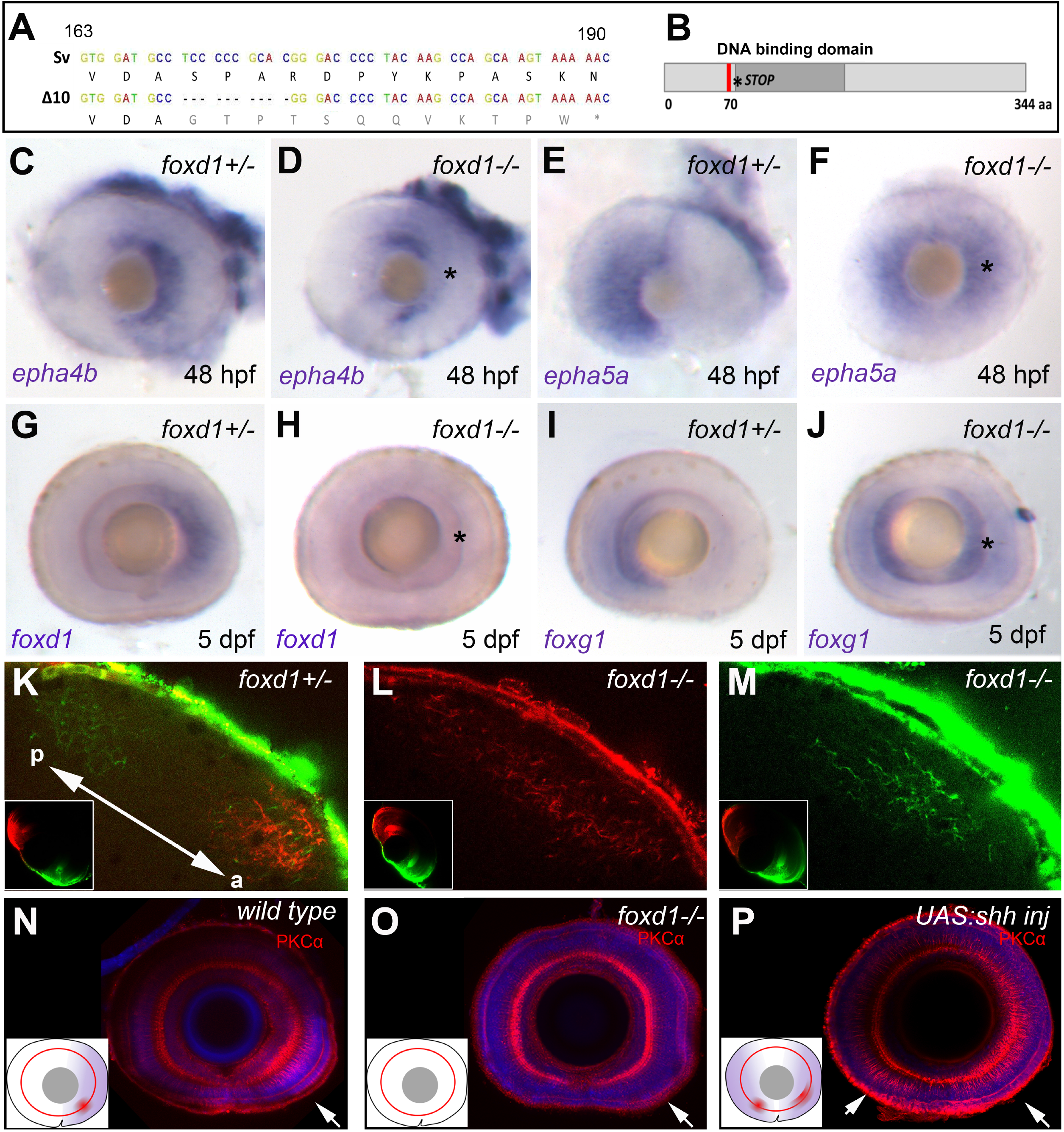
Loss of *foxd1* results in retinae with expanded nasal, and reduced temporal character. (A) *foxd1* sequence comparison between nucleotides 163 and a 190 of the open reading frame, highlighting the 10 base-pair deletion (Δ10, bottom row) in the *foxd1*^*cbm16*^ mutant. (B) Diagrammatic representation of the Foxd1 protein, depicting in red the deleted region in the *cbm16* allele. The asterisk in (A-B) marks the position at which a newly generated stop codon is present. (C-J) Lateral views with nasal to the left of 48hpf (C-F) and 5dpf (G-J) retinae showing expression of nasal or temporal markers in *foxd1*^*−/−*^ mutants and siblings. The markers assessed and the genotypes of the retinae are detailed bottom left and top right respectively. Asterisk in (D,F,H,J) highlights altered expression in *foxd1* mutants. (K-M) Dorsal view of the tectum of *foxd1*^*−/−*^ mutant (L-M) and sibling embryos (K), with nasal RGC arbors labelled with DiO (green) and temporal RGC arbors with DiI (red). Insets in (K-M) show dorsal views of the corresponding eye. Tectum and eye orientation in panels (K-M) is indicated by the double arrow in (K). (N-P) PKC α immunostaining (red) in lateral views with nasal to the left of 8dpf wildtype (N), *foxd1*^*−/−*^ mutant (O) and *tg{rx3:Gal4};UAS:Shh* (P) eyes. Nuclei are counterstained with dapi (blue). Insets in (N-P) are schematics of the eyes in the corresponding panels, highlighting the extent of temporal character (purple), as assessed by gene expression, and the areas of enriched PKC α (red). Arrows indicate the position of PKC α enrichment.

*foxd1*^*−/−*^ embryos developed at least up to 9 dpf, showing no apparent morphological malformations in the retina or elsewhere (data not shown). However, the temporal retinal marker *epha4b*, and *foxd1* itself, were downregulated in the mutant eyes (**Figure 2C-D,G-H**). This downregulation was accompanied by a mirror-image duplication of expression of the nasal retinal markers *foxg1* and *ephA5a* in the temporal half of the retina (**Figure 2E-F, I-J** and **Figure S3A-D**). Of note, downregulation of *foxd1* expression in *foxd1*^*−/−*^ embryos was not due to non-sense mediated decay, since expression in other regions of the embryo was not affected (**Figure S4**). The retinal phenotype of *foxd1*^*−/−*^ mutants was phenocopied in F0 “crispants” generated by simultaneous injection of three guides targeting different regions of the *foxd1* locus (**Figure S3G**; Kroll et al., 2021; Materials and Methods). All genotyped crispant embryos showed gene editing (n=16; **Figure S3H**) and over 90% of the analysed embryos (65 out of 72) showed the expansion in *foxg1* expression characteristic of *foxd1* mutants (**Figure S3E-F**).

Analysis of retinotectal projections revealed an aberrant pattern of tectal innervation in *foxd1*^*−/−*^ mutants. In control conditions, temporally located RGCs, injected with DiI, innervated the anterior portion of the contralateral tectum, while nasally located RGCs, injected with DiO, innervated the posterior tectum (**Figure 2K**; inset shows the corresponding eye, n=10 eyes). In *foxd1*^*−/−*^ mutants, RGC terminals (labelled as for control conditions) from both nasal or temporal halves of the retina showed overlapping arborisations across much of the tectum (**Figure 2L-M**; insets show the corresponding eye, n=12 eyes). This result indicates that in the absence of RGCs with temporal character, axons from RGCs with nasal character expand their arbours throughout the tectum (Suetterlin et al., 2012). Overall, these results suggest that the *foxd1*^*−/−*^ mutant retina has all-nasal character.

### HAA specification is linked to naso-temporal patterning

In fish, the ventro-temporal region of the retina (also called the *area temporalis*; Schmitt and Dowling, 1999) bears several structural specialisations that are thought to provide high visual acuity and support prey detection: a high density of U-cones (invested in detecting light in the ultraviolet spectrum) and low density of rods (Yoshimatsu et al., 2020; Zimmermann et al., 2018), a specialised inner retina neuronal circuit (Mangrum et al., 2002; Pita et al., 2015) and a characteristic number, shape and position of synaptic terminals at the inner plexiform layer (Zhou et al., 2020; Zimmermann et al., 2018). Immunostaining with anti-PKCα highlights the cell anisotropies found in the inner nuclear layer and inner plexiform layer at the level of the HAA (Zimmermann et al., 2018).

In contrast to wildtype zebrafish, where PKCα immunostaining was enriched in the ventro-temporal region of the retina (**Figure 2N**), PKCα enrichment was lost in *foxd1*^*−/−*^ retinae (**Figure 2O**), suggesting that HAA-specific features are absent in *foxd1*^*−/−*^ mutants. Despite this absence of HAA-specific features, *foxd1*^*−/−*^ retinae showed an otherwise normal PKCα expression and architecture (**Figure 2** and not shown).

To determine whether changes in HAA markers are disrupted in other conditions that affect naso-temporal patterning, we analysed the opposite condition, in which embryos present a retina with all-temporal character. To generate this condition, we misexpressed Shh throughout the developing eye using the Gal4/UAS system (Distel et al., 2010; Halpern et al., 2008; Paquet et al., 2009) as previously described (Hernández-Bejarano et al., 2015). As shown in a previous report (Hernández-Bejarano et al., 2015), *Tg{rx3:Gal4};UAS:Shh* embryos showed an expansion of *foxd1* expression throughout the retina at the expense of *foxg1*.

*Tg{rx3:Gal4};UAS:Shh* retinae showed a dorso-temporal expansion in the HAA-associated enrichment of PKCα (**Figure 2P**, arrows; compare with wild type in **Figure 2N**), suggesting that the HAA was not only present, but expanded in this condition. Notably, an ectopic patch of PKCα enrichment also appeared to be present in the ventro-nasal half of the retina (**Figure 2P**). Overall, these results suggest that the establishment of nasal and temporal character is linked to the formation of retinal specialisations such as the HAA.

### Visual function is impaired in *foxd1* mutants

Perturbation of naso-temporal patterning and retinotectal projections in *foxd1*^*−/−*^ mutants is likely to affect visual function. We therefore examined two robust visually guided stabilisation reflexes elicited by whole-field motion: the optokinetic and the optomotor responses (OKR and OMR, respectively). The OKR is a visually driven compensatory eye movement that minimises retinal image slip to stabilise gaze. During OKR, animals perform slow phase eye rotations in the direction of visual motion with intermittent fast phase saccades to reset eye position. In the OMR, zebrafish larvae turn and swim in the direction of perceived whole field motion (Neuhauss et al., 1999).

*foxd1*^*−/−*^ mutants were substantially impaired in OKR assays. We presented 6 dpf tethered larvae with rotating visual gratings and quantified the gain (ratio of eye velocity to stimulus velocity) of the slow phase of the optokinetic nystagmus as a function of stimulus spatial frequency, temporal frequency and contrast. Gain was reduced by around 80% in all conditions in *foxd1*^*−/−*^ mutants (**Figure 3B**, red) as compared to wild types and heterozygote siblings (**Figure 3B**, black/blue). While this could reflect deficits in visual perception, we also observed a reduction in oculomotor range and in peak eye velocity (**Figure 3D**) suggesting that oculomotor defects might contribute to the phenotype. We therefore tested a second stabilisation reflex evoked by whole field visual motion, and measured OMR performance. Whereas wildtype larvae robustly turned in the direction of whole-field motion, *foxd1* crispant larvae were more prone to make mistakes and only performed about 67% turns in the correct direction (**Figure 3F**). *foxd1*crispant larvae did not show evidence of locomotor dysfunction as they performed a similar total number of swim bouts compared to wild types and displayed a normal range of turn angles (**Figure 3G and Figure S5**). Taken together, these data indicate that loss of FoxD1 function leads to impaired visual motion perception.

**Figure 3:**
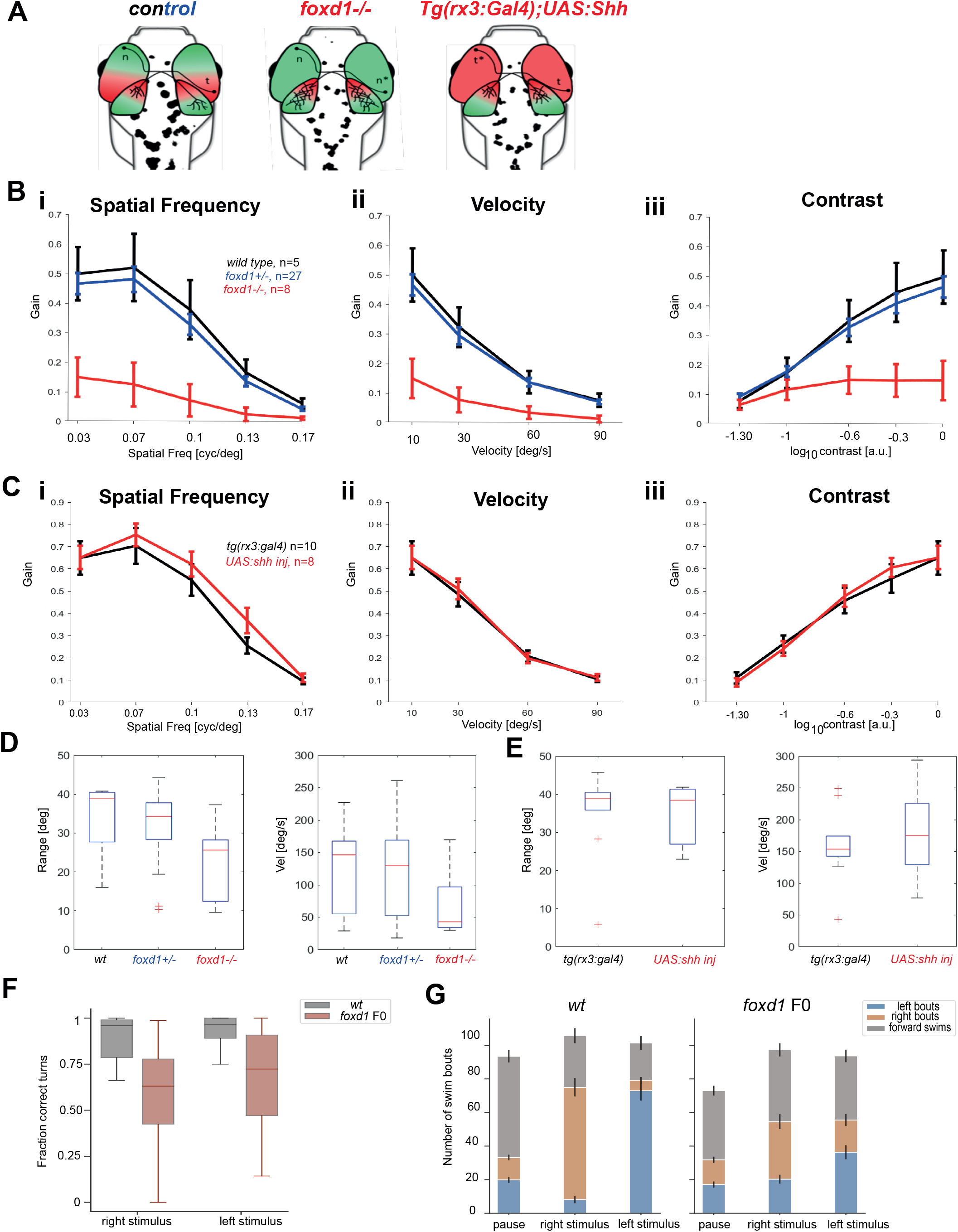
OKR assays reveal severe visual deficiencies in *foxd1*^*−/−*^ but not *tg{rx3:Gal4};UAS:Shh* embryos. (A) Diagrammatic representation of the changes in naso-temporal pattern and retinotectal projections observed in the four categories of fish tested by OKR. Genotype is detailed at the top of each diagram. (B-C) Graphs detailing eye gain (expressed in a 0-1 range) in response to changes in spatial frequency (expressed in cycles per screen; Bi,Ci), grating velocity (expressed in degrees per second; Bii,Cii) and contrast (expressed in a 0-1 range; Biii,Ciii) in wild type (black, n=5 in B), *foxd1*^*+/−*^ (blue, n=27 in B), *foxd1*^*−/−*^ (red, n=8; in B), *tg{rx3:Gal4}* (black [rx3:gal4], n=10 in C) and *tg{rx3:Gal4};UAS:Shh* (red, [+UAS:shh], n=8 in C). Graphical representation was generated with GraphPad Prism5 (ANOVA double test, followed by a Bonferroni test) and statistical significance calculated with a t-student test (***p<0.001). (D-E) Range (expressed in degrees) of slow phase eye movement (left graph) and peak velocity (expressed in degrees per second) during fast phase eye movement (right graph) in *foxd1*^*−/−*^ versus wildtype and *foxd1*^*+/−*^ fish (D), and *tg{rx3:Gal4}* versus *tg{rx3:Gal4};UAS:Shh* fish (E). (F) Fraction of correct turns for wildtype larvae (n=21) and *foxd1* crispants (*foxd1 F0*, n=21) when subjected to left- and right-oriented whole field motion stimuli. Only directional bouts (i.e. left- and rightward swims, without forward swims) were considered for quantification. (G) Number of swim bouts during pause intervals and whole field motion stimuli for wildtype larvae (left, n=21) and *foxd1* crispants (right, n=21). Bar heights represent the means across all fish in each group; error bars represent the standard error of the mean (SEM).

To determine whether impairment in visual motion perception is characteristic of other conditions that affect naso-temporal patterning and retinotectal projections, we assessed visual performance in *Tg{rx3:Gal4};UAS:Shh* fish with double temporal retinae. RGCs positioned in the nasal region of *Tg{rx3:Gal4};UAS:Shh* embryos projected more anteriorly in the tectum, overlapping with the projections from temporally located RGCs (**Figure 3A**; Hernández-Bejarano et al., 2015). Despite altered naso-temporal character and retinotectal projections, we did not observe any deficits in OKR performance in *Tg{rx3:Gal4};UAS:Shh* larvae (**Figure 3C**, red; compare with uninjected *Tg{rx3:Gal4}* larvae in black). These results suggest that the difference in OKR performance in *foxd1*^*−/−*^ mutants and *Tg{rx3:Gal4};UAS:Shh* embryos is associated with the absence or presence, respectively, of temporal retinal character and, potentially, associated to connectivity alterations.

## Discussion

In this study we identify the eye field specification gene *rx3* as a regulator of *foxd1* expression, thereby further resolving the transcriptional and signalling pathways that lead to naso-temporal retinal pattern (**Figure 4**). We propose that *rx3* expression in the nascent eye field confers to this whole domain competence to express *foxd1. foxd1* is repressed in the dorsal half of the eye primordium by Fgf activity and promoted in the ventral half by Hh activity (this study and Hernández-Bejarano et al., 2015). Fgf activity also promotes the expression of the nasal determinant *foxg1* and FoxG1 and FoxD1 subsequently engage in a negative cross-regulatory relationship that refines and maintains the border between nasal and temporal retinal domains (Hernández-Bejarano et al., 2015).

**Figure 4:**
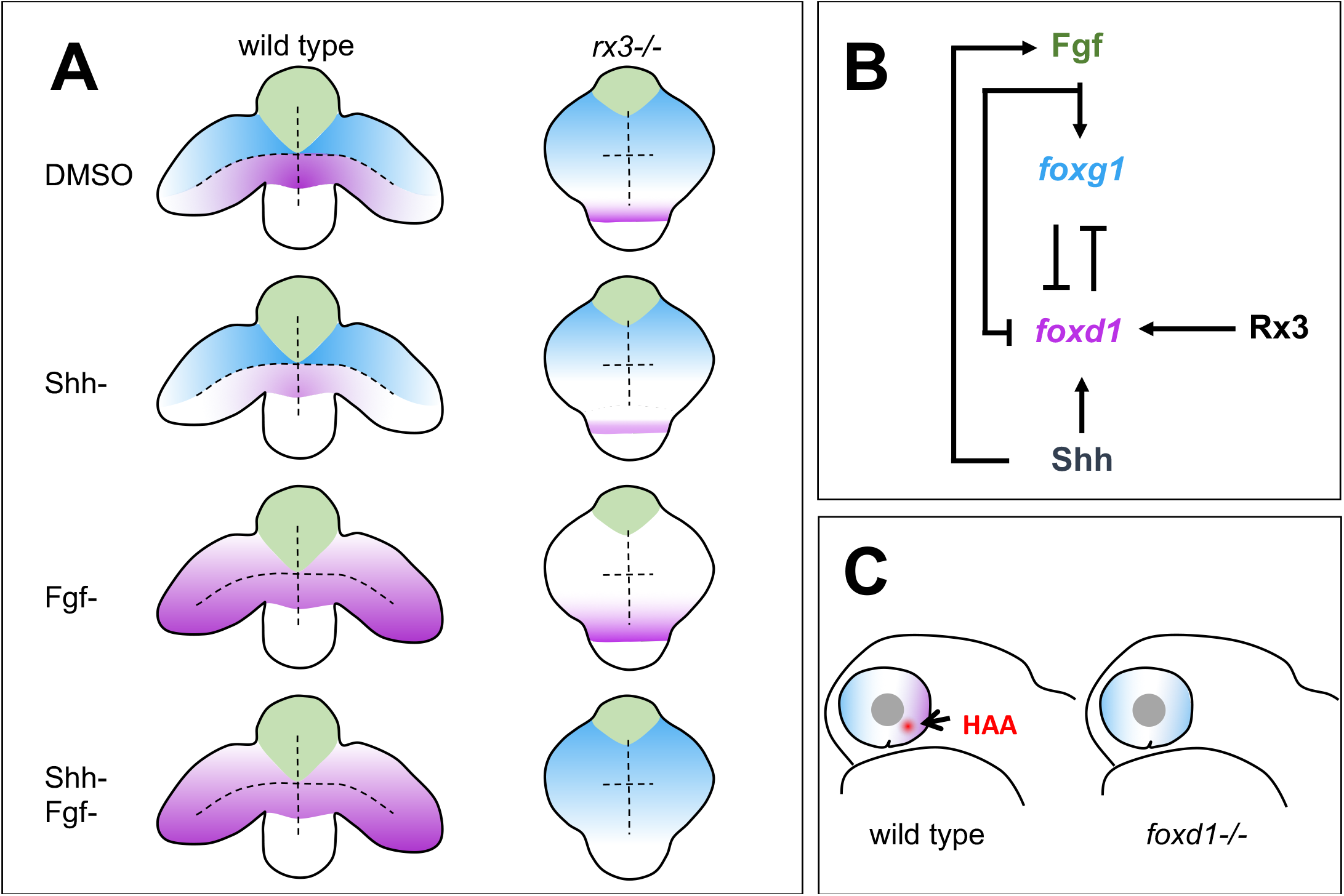
Summary of roles for Rx3, Fgfs and Shh in establishment of nasal and temporal character in the developing eye primordium. (A) Schematic representation of wildtype and *rx3*^*−/−*^ embryos depicting the changes in *foxg1* (blue) and *foxd1* (magenta) expression upon abrogation of Shh and Fgf activities, singly or in combination. (B) Diagram detailing the proposed regulatory network controlling NT patterning. Shh promotes *foxd1* expression in the ventral (future temporal) half of the optic primordium. Fgfs induce *foxg1* in the dorsal (future nasal) optic primordium and repress *foxd1* expression, contributing to its confinement ventrally. Cross-repression between FoxG1 and FoxD1 at the border between the dorsal and ventral halves of the eye primordium subsequently refines the naso-temporal subdivision. Although induction of *shh* and *fgf8* expression occurs independently, *fgf8* expression is lower in absence of Shh signalling (Hernández-Bejarano et al., 2015) suggesting Hh signalling initially promotes Fgf signalling. Rx3 induces *foxd1* expression and provides competence to optic vesicle cells to enhance *foxd1* expression in prospective temporal retina in response to Shh signalling. (C) Schematic representation of a differentiated optic cup with nasal (blue) and temporal (magenta) character, and the HAA (red) highlighted. Nasal character is expanded and HAA and temporal character are absent in *foxd1*^*−/−*^ mutants (right) as compared to the wild type (left).

The requirement of Rx3 to promote *foxd1* expression is most clearly revealed when the activity of other regulators is removed. In the absence of Fgf and Shh, *foxd1* expands throughout the optic vesicles. Removing Rx3 activity in this condition leads to loss of *foxd1* expression with a concomitant expansion of *foxg1* expression, confirming the requirement for Rx3 in the induction of *foxd1* expression.

An *area temporalis* in the zebrafish retina characterized by tightly packed cones was identified more than 20 years ago (Schmitt and Dowling, 1999). This area corresponds to the HAA, and more recent studies have shown that it bears a distinctive density of particular neuronal types and a specialised neuronal circuitry (Mangrum et al., 2002; Pita et al., 2015; Yoshimatsu et al., 2020; Zhou et al., 2020; Zimmermann et al., 2018). Our analysis suggests that the establishment of temporal retinal character is required for HAA formation. *foxd1*^*−/−*^ mutants show an expansion of nasal retinal markers at the expense of temporal markers. Accumulation of PKCα in the ventro-temporal retina, which highlights the structural features of the HAA inner retina and inner plexiform layer (Zimmermann et al., 2018), is lost in *foxd1*^*−/−*^ mutants, suggesting that the mutant retina is devoid of at least some HAA features.

Not only the HAA, but also other structural anisotropies in the temporal retina may be affected in *foxd1*^*−/−*^ mutants. Recent studies suggest that the OKR is driven mainly by stimuli covering the central visual field (Dehmelt et al., 2021) and predict that the HAA specialisations in the temporal retina are dispensable for OKR. However, OKR and OMR tests in *foxd1*^*−/−*^ mutants reveal defective whole-field motion perception in these larvae. This suggests that either the HAA or other regions of temporal retina required for this visual response, are affected in *foxd1*^*−/−*^ mutants. Conversely, larvae with temporal character expanded into the nasal domain (*Tg{rx3:Gal4};UAS:Shh*) show an overtly normal OKR. These data suggest that perturbation of nasal retinal character does not interfere with this aspect of visual function, and supports the idea that at least some aspects of the OKR are predominantly driven by retina with temporal character.

Ablation and lesion experiments in both fish and other vertebrate models suggest that the execution of the OKR is driven mainly by the pretectum and can occur in the absence of an intact tectum (Flandrin and Jeannerod, 1981; Roeser and Baier, 2003). Here we show that defective whole-field motion perception in *foxd1*^*−/−*^ mutants, which show extensive disruption of retinotectal projections, is perturbed, but our analysis does not allow us to determine whether these two phenotypes are causally related as innervation of other retinorecipient areas may also be perturbed in *foxd1*^*−/−*^ mutants. Indeed, some pretectal arborisation fields are preferentially innervated by the temporal retina (Robles et al., 2014), and our analysis did not determine whether these are affected in *foxd1*^*−/−*^ mutants.

In summary, our study uncovers a mechanistic link between early naso-temporal patterning events in the eye primordium and the establishment of functionally distinct regions in the retina. Despite the importance of naso-temporal patterning for accurate RGC connectivity to specific retino-recipient areas, analysis of the visual function in *foxd1*^*−/−*^ mutants and *Tg{rx3:Gal4};UAS:Shh* fish reveals a surprising robustness in the ability of the larvae to perceive stimuli in the absence of nasal or temporal-specific retinal character. These lines of fish provide an exciting new avenue to further our understanding of the role of nasal and temporal retinal character in mediating specific visual functions.

## Materials and Methods

### Fish lines and husbandry

*AB* and *tupl* wildtype zebrafish strains, the transgenic line *Tg{rx3::Gal4-VP16}*^*vu271Tg*^ (Weiss et al., 2012), and mutant lines *chk*^*ne2611*^ (Loosli et al., 2003) and *foxd1*^*cbm16*^ were maintained and bred according to standard procedures (Aleström et al., 2020; Westerfield, 1993). All experiments conform to the guidelines from the European Community Directive and the British (Animal Scientific Procedures Act 1986) and Spanish (Real Decreto 53/2013) legislation for the experimental use of animals.

### Generation of the foxd1 mutant

The sequence to target in the *foxd1* open reading frame (ENSDARG00000029179) was selected using the UCSC Genome Browser (http://genome.uscs.edu). A DNA oligonucleotide bearing the target sequence (GCTTGTAGGGGTCCCGTGC; positions 435-417 on the reverse strand of the transcript), the T7 RNA polymerase binding site and the sequence complementary to the oligoB universal primer was commercially obtained, expanded with Expand High Fidelity DNA polymerase (Roche) and transcribed with the Maxi script T7 kit (NEB), following manufacturers’ instructions. Cas9 mRNA was generated by linearisation and transcription from the PCS2-nCas9n (Addgene, #47929) clone, purified (PCR cleanup kit, Roche) and transcribed (SP6 mMessage mMachine Kit Ambion), following manufacturers’ instructions. F0 founders were generated by co-injection of guide RNA (25ng/μl) and Cas9 mRNA (300ng/μl) in *AB/tupl* embryos at one cell stage. Cleavage efficiency was assessed in pools of injected embryos by CRISPR-STAT analysis (Carrington et al., 2015). Genotyping of F0 founders and their progeny was performed by CRISPR-STAT or HRM analysis from genomic DNA samples obtained from tail fin biopsies. The primers used are detailed in **Table S1**.

### Generation of multi-guide foxd1 crispants

The phenotype of the *foxd1*^*cbm16*^ mutants was phenocopied by multi-guide injection, following the protocol previously published by (Kroll et al., 2021). Three synthetic RNA guides were designed (1AA, 1AB and 1AC guides, Supplementary Table 1) and ordered to Integrated DNA Technologies (IDT). Guides were annealed to the tracrRNA oligonucleotide (IDT#1072532), assembled with Cas9 protein (IDT#1081058) and injected into one-cell stage wildtype embryos, as previously described (Kroll et al., 2021). A subset of the injected embryos was genotyped by HRMA using the HRMA primers described in **Table S1**, to confirm the presence of gene editing.

### Microinjection and drug treatments

Treatments with SU5402 and cyclopamine were performed as previously described (Hernández-Bejarano et al., 2015). SU5402 (Calbiochem) and cyclopamine (Calbiochem) were applied at a concentration of 10μM and 100μM, respectively, to pools of embryos derived from the mating of *rx3*^*+/−*^ parental fish, at the corresponding stage of embryonic development. 30-50 embryos were treated and processed together per experiment/marker. *rx3*^*−/−*^ embryos were recovered at the expected mendelian proportions and the phenotypes observed were fully penetrant.

Overexpression of *shh* in the optic vesicle under the control of the Gal4/UAS system (Halpern et al., 2008) was performed by injecting 20-30ng of bidirectional GFP:UAS:*shh* plasmid DNA into the cell of one-cell stage *Tg{rx3::Gal4-VP16}*^*vu271Tg*^ embryos. Only embryos with homogeneous GFP expression in the optic vesicles were selected and processed for analysis (as described in Hernández-Bejarano et al., 2015).

### mRNA detection and immunolabelling

mRNA detection (preparation of RNA antisense probes and whole mount *in situ* hybridisation) was performed as previously described (Hernández-Bejarano et al., 2015). Immunolabelling of 8 dpf wildtype, *foxd1* and *Tg{rx3::Gal4};*UAS:*Shh* retinae with anti-PKCα (Sigma, P4334, 1:100), Alexa-coupled secondary antibodies (Jackson ImmunoResearch, 1:500) and the nuclear dye DAPI was performed on RPE-dissected retinae as described in (Zimmermann et al., 2018).

### Tracing of retinotectal projections

Nasal and temporal RGCs were labelled with DiO and DiI, respectively in 6 dpf *wild type, foxd1*^*−/−*^ and *Tg{rx3:Gal4};*UAS:*Shh* retinae previously fixed in 4% paraformaldehyde. Labelled larvae were incubated at room temperature for at least 24 hours before imaging. Each tectum and its corresponding eye were sequentially imaged (Hernández-Bejarano et al., 2015).

### Optokinetic response test (OKR)

6 dpf larvae were anesthetised with tricaine (MS-222, Sigma), embedded in 1% low-melting point agarose (Sigma) and immersed in embryo medium. A small portion of agarose was removed from around the eyes to allow movement. Once larvae had recovered from anaesthesia, OKR performance was tested by presenting rotating visual gratings that varied in size, contrast and speed. We presented a set of 12 unique stimuli that was repeated twice for each animal. Horizontal eye movement was tracked under infrared illumination (850 nm) with an AVT Pike camera (100 frames per second). Stimulus presentation and machine vision were controlled using LabVIEW (National Instruments). Slow phase gain, saccadic eye velocity and oculomotor range were computed using custom MATLAB software (https://IsaacBianco@bitbucket.org/biancolab/okrsuite.git). *foxd1*^*−/−*^ mutants (n=8) were compared to wildtype (n=5) and *foxd1*^*+/−*^ heterozygotes (n=27) and *Tg{rx3:Gal4};*UAS:*Shh* (n=8) were compared to *Tg{rx3:Gal4}* (n=10) uninjected larvae. Significance was calculated with a two-way ANOVA test, followed by a Bonferroni test (GraphPad Prism5). Graphical representation shows eye gain (0-1 range, *y* axis) at variable speed (cycles per screen), frequency (degrees per second) or contrast (0-1 range; *x* axis).

### Optomotor response assay (OMR)

6 dpf zebrafish larvae were individually placed in a 6 cm Petri dish and presented from below via a projector with moving grating stimuli in a closed loop manner. The customized stimulus protocol and tracking of the freely moving fish was implemented using the Stytra software package (Štih et al., 2019). The time course of the assay consisted of 5 repetitions of rightwards and leftwards moving gratings (20 seconds each), separated by 10 seconds of static gratings (pause), for a total time of 5 minutes per fish. 21 wildtype and 21 *foxd1*-crispant larvae were subjected to the OMR assay. Custom behavioral analysis was implemented using Python and the bouter package (Stih et al., 2022).

### Imaging and data processing

DiI/DiO-traced embryos were embedded in low melting point agarose (Sigma) at 1-1.5% in PBS for confocal imaging using a 40X (0.8NA) long-working distance water immersion lens. A Zeiss LSM710 confocal microscopy system was used for image acquisition. Immunolabelled 8dpf retinae were imaged using a Leica SP8 microscope with at 25x water immersion lens.

*In situ* hybridised embryos and dissected eyes were mounted flat in a drop of glycerol and dorsal images were acquired with a 20X (0.70NA) dry lens using a Leica CTR 5000 microscope connected to a digital camera (Leica DFC 500), and operated by Leica software. Some embryos were embedded in gelatine/BSA for vibratome sectioning as previously described (Hernández-Bejarano et al., 2015; Sánchez-Arrones et al., 2013). Alternatively, embryos were cryoprotected in sucrose 30% and embedded in OCT (Sakura Fintek) for cryo-sectioning as previously described (Cavodeassi et al., 2013). Sections (16 μm) were obtained using a Leica VT1000S vibratome or a Leica cryostat, mounted in glycerol, and imaged with a 40X (1.3 NA) oil-immersion lens. Images in Supplementary Figure 3 were acquired using a 20X dry lens using a Nikon Eclipse microscope connected to a digital camera (DS-Fi3) and operated by Nikon software (NIS-Elements).

Raw confocal images were processed with ImageJ. Processed images were exported as TIFF files and all figures were composed using Photoshop.

## Acknowledgements

We would like to thank Alexander Picker, discussions with whom inspired and prompted this study; and Kenzo Ivanovitch, Paride Antinucci and Paris Ataliotis for reading and providing insightful feedback during the writing of this manuscript. The animal facility at the CBMSO, the zebrafish facility at UCL and the Biological Research Facility at St. Georges, University of London (SGUL) are acknowledged for their help with animal care. The imaging facilities at the CBMSO, UCL and SGUL are also acknowledged for their help with image acquisition.

