## Supplementary information for "Foxd1 dependent induction of temporal retinal character is required for visual function"

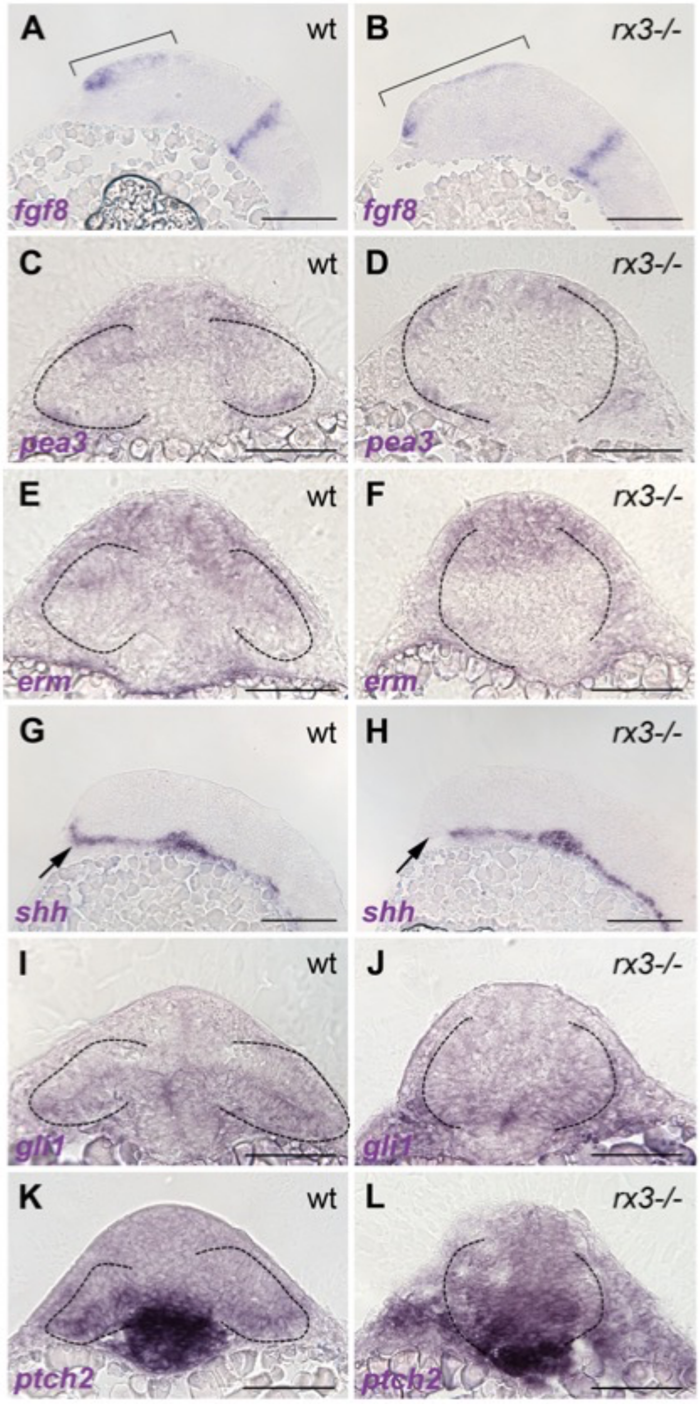


***Supplementary figure 1:*** **expression of Fgf and Shh pathway genes is similar in** ***$rx3^{-/-}$*** **mutants and siblings.**

**(**A-L) Sagittal (A,B; G,H) and frontal sections at the level of the forming telencephalon, optic vesicles and hypothalamus (C-F;I-L), with dorsal up. Genes analysed and genotypes are shown bottom left and top right respectively. Brackets in (A-B) highlight the extent of *fgf8* expression in the prospective telencephalon; arrows in (G-H) highlight the extent of *shh* expression in the anterior hypothalamus. All embryos are 10ss.

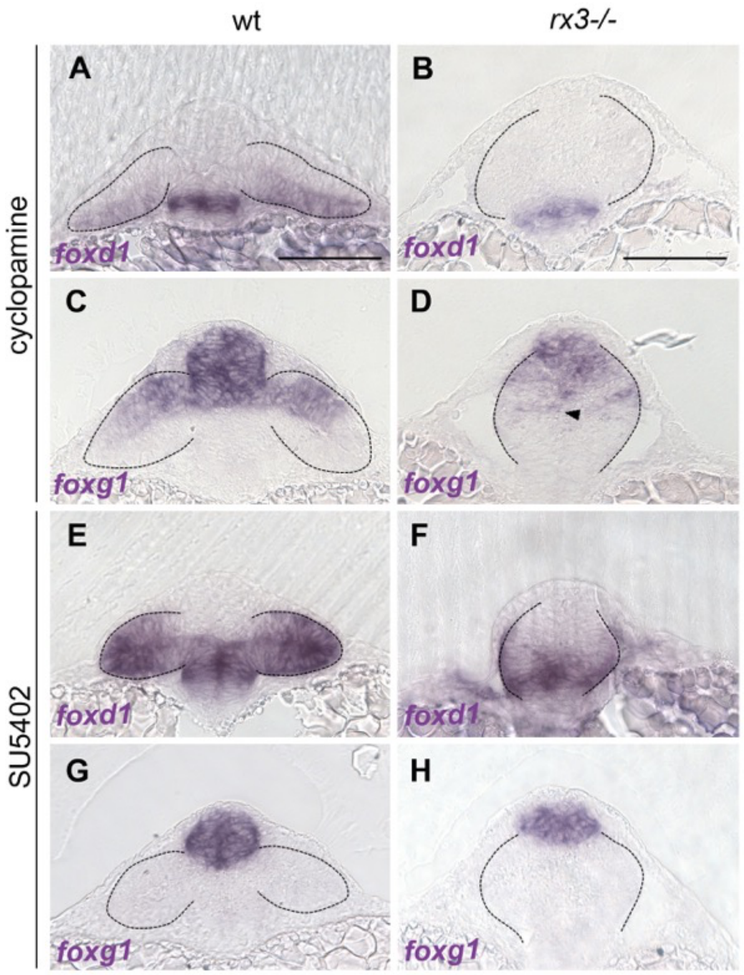


***Supplementary figure 2:*** **abrogation of Shh or Fgf activity in *rx3* mutant embryos.**

Frontal sections at the level of the forming telencephalon, optic vesicles and hypothalamus with dorsal up. Genes analysed and genotypes are shown bottom left and top respectively.

(A-D) Cyclopamine treated 10ss *rx3*$^{-/-}$ (B,D) and sibling embryos (A,C).

(E-H) SU5402 treated 10ss *rx3*$^{-/-}$ (F,H) and sibling embryos (E,G).

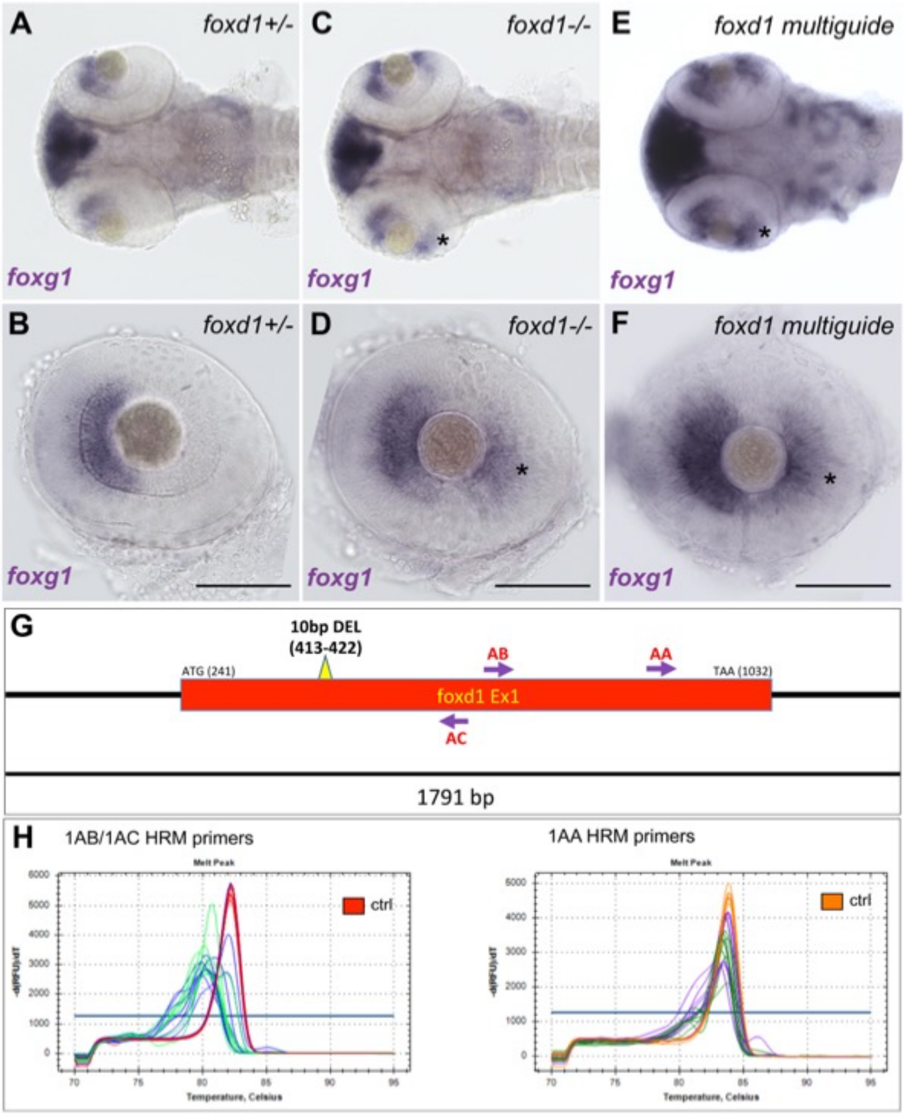


***Supplementary figure 3: foxd1* "crispants" reproduce the *foxg1* expression expansion phenotype observed in *foxd1* mutants.**

(A-F) dorsal views of brains and eyes (A,C,E) and lateral views of eyes (B,D,F) showing *foxg1* expression in 60-72hpf wildtype (A-B), *foxd1*$^{-/-}$ mutants (C-D) and *foxd1* crispants (E-F). Note the expansion of *foxg1* to the temporal retina (asterisk in C-F).
(G) Schematic of the *foxd1* locus with the regions targeted by the CRISPR guides highlighted.
(H) HRM profile in crispant embryos as compared to the wildtype profile. Two sets of primers were used to assess *indels* induced by guide 1AA (right panel) and guides 1AB/1AC (left panel). The red profile on the left panel and orange profile on the right panel correspond to wildtype controls. All genotyped embryos showed *indels* in both regions.

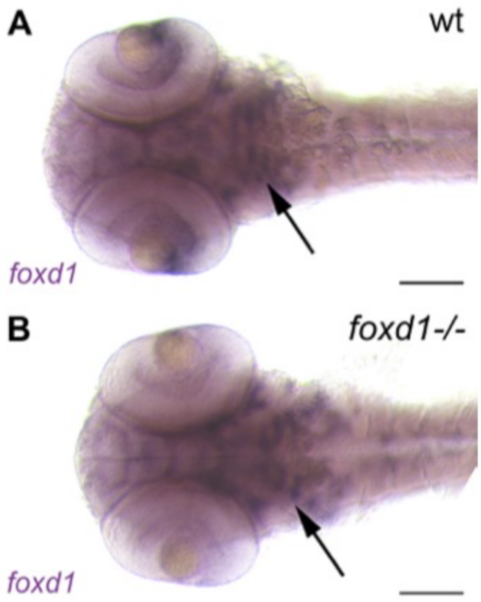


***Supplementary figure 4:*** **loss of *foxd1* expression in the temporal retina of *foxd1* mutants is not due to nonsense-mediated RNA decay.**

Ventral views of heads showing *foxg1* expression in 60-72hpf wildtype (A) and *foxd1*$^{-/-}$ mutants (B). Note *foxd1* expression in the brain and branchial arches regions (arrows) is maintained in *foxd1*$^{-/-}$ mutants (B, compare to A), despite being absent in the retina. Anterior is to the left.

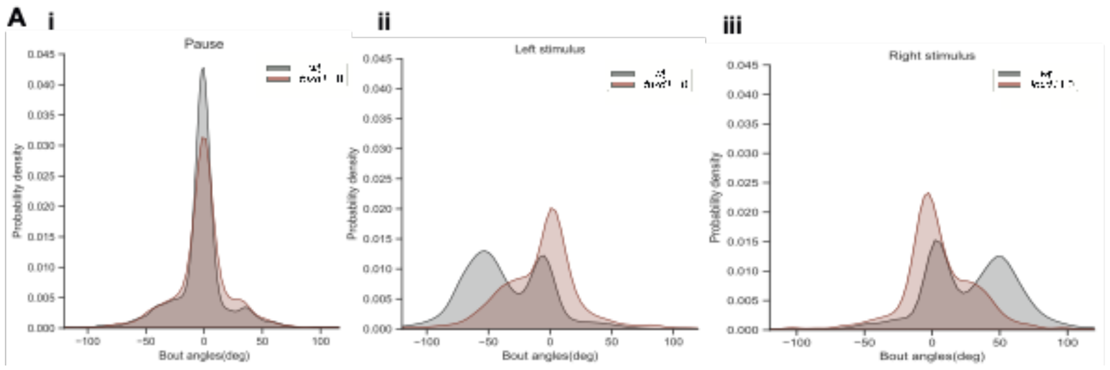


***Supplementary Figure 5:* range of swim bout angles is similar in wildtype and *foxd1* larvae.**

Probability densities of swim bout angles for wildtype and *foxd1* crispant larvae, during the pause interval (i), the leftwards (ii) and rightwards (iii) oriented whole field motion stimulus.

***Supplementary table 1: Crispr guides and genotyping primers***

| Primers used | Sequence |
|---|---|
| foxd1 original CRISPR guide | GCTTGTAGGGGTCCCGTGC |
| foxd1 original CRISPR-STAT-fwd | TGTAAAACGACGGCCAGTTCAGATGCACGACGAGATC |
| foxd1 original CRISPR-STAT-rev | GTGTCTTGTCACAAATCTCGCTCAGC |
| foxd1 original CRISPR HRM-fwd | CAGATGCACGACGAGATCCT |
| foxd1 original CRISPR HRM-rev | GAAGTCACAAATCTCGCTCAGC |
| foxd1.1AA guide | GGCTATGGACCCTACGGTTG |
| foxd1.1AB guide | TCAAGATACCACGAGAGCCC |
| foxd1.1AC guide | CGGATAGAGTTCTGCCAAGC |
| foxd1-1AB/1AC HRM-fwd | GCTGAGCGAGATTTGTGACTTC |
| foxd1-1AB/1AC HRM-rev | CTAGCGTCCAGTAGTTGCCTTTG |
| foxd1-1AA HRM-fwd | GCGAGCACGGAGGTTTTCTTCC |
| foxd1-1AA HRM-rev | GAAAGGCGAGGAGCGCGGAATG |
